# Deep learning and host variable embedding augment microbiome-based simultaneous detection of multiple diseases

**DOI:** 10.1101/2023.05.16.541058

**Authors:** Shunyao Wu, Zhiruo Li, Yuzhu Chen, Mingqian Zhang, Yangyang Sun, Jieqi Xing, Fengyang Zhao, Shi Huang, Rob Knight, Xiaoquan Su

## Abstract

Microbiome has emerged as a promising indicator or predictor of human diseases. However, previous studies typically labeled each specimen as either healthy or with a specific disease, ignoring the prevalence of complications or comorbidities in actual cohorts, which may confound the microbial-disease associations. For instance, a patient may suffer from multiple diseases, making it challenging to detect their health status accurately. Furthermore, host phenotypes such as physiological characteristics and lifestyles can alter the microbiome structure, but this information has not yet been fully utilized in data models. To address these issues, we propose a highly explainable deep learning (DL) method called Meta-Spec. Using a deep neural network (DNN) based approach, it encodes and embeds the refined host variables with microbiome features, enabling the detection of multiple diseases and their correlations simultaneously. Our experiments showed that Meta-Spec outperforms regular machine learning (ML) strategies for multi-label disease screening in several cohorts. More importantly, Meta-Spec can successfully detect comorbidities that are often missed by regular ML approaches. In addition, due to its high interpretability, Meta-Spec captures key factors that shape disease patterns from host variables and microbial members. Hence, these efforts improve the feasibility and sensitivity of microbiome-based disease screening in practical scenarios, representing a significant step towards personalized medicine and better health outcomes.

## Introduction

The dynamics of the human microbiome are closely associated with numerous diseases [1, 2]. In recent years, the increasing volume and diversity of microbiome data have led to the widespread adoption of machine learning (ML) based approaches for disease detection and recognition [3-5]. Typically, ML classifiers take taxonomic or functional features of microbiomes in different health conditions for training, and construct classification models to predict the status of new samples. In this scenario, microbiome cohorts for research are always well-designed that each specimen is marked with only one definitive status, denoted as “label”, i.e., either healthy or with a specific disease [6, 7]. Such effort reduces the confounding of complex factors in the experimental design and makes ‘single-label classification’ a typical strategy (**Figure 1a**). However, complications or comorbidities are prevalent in actual cohorts (**Figure 1b**). For example, in the American Gut Project (AGP) [8], ∼61% of patients were diagnosed with at least two diseases. Our recent study has shown that gut microbiomes with comorbidities can have distinct microbial patterns from those with a single disease, even though they share common biomarkers [9]. As a result, comorbidities can significantly disrupt disease detection.

**Figure 1.**
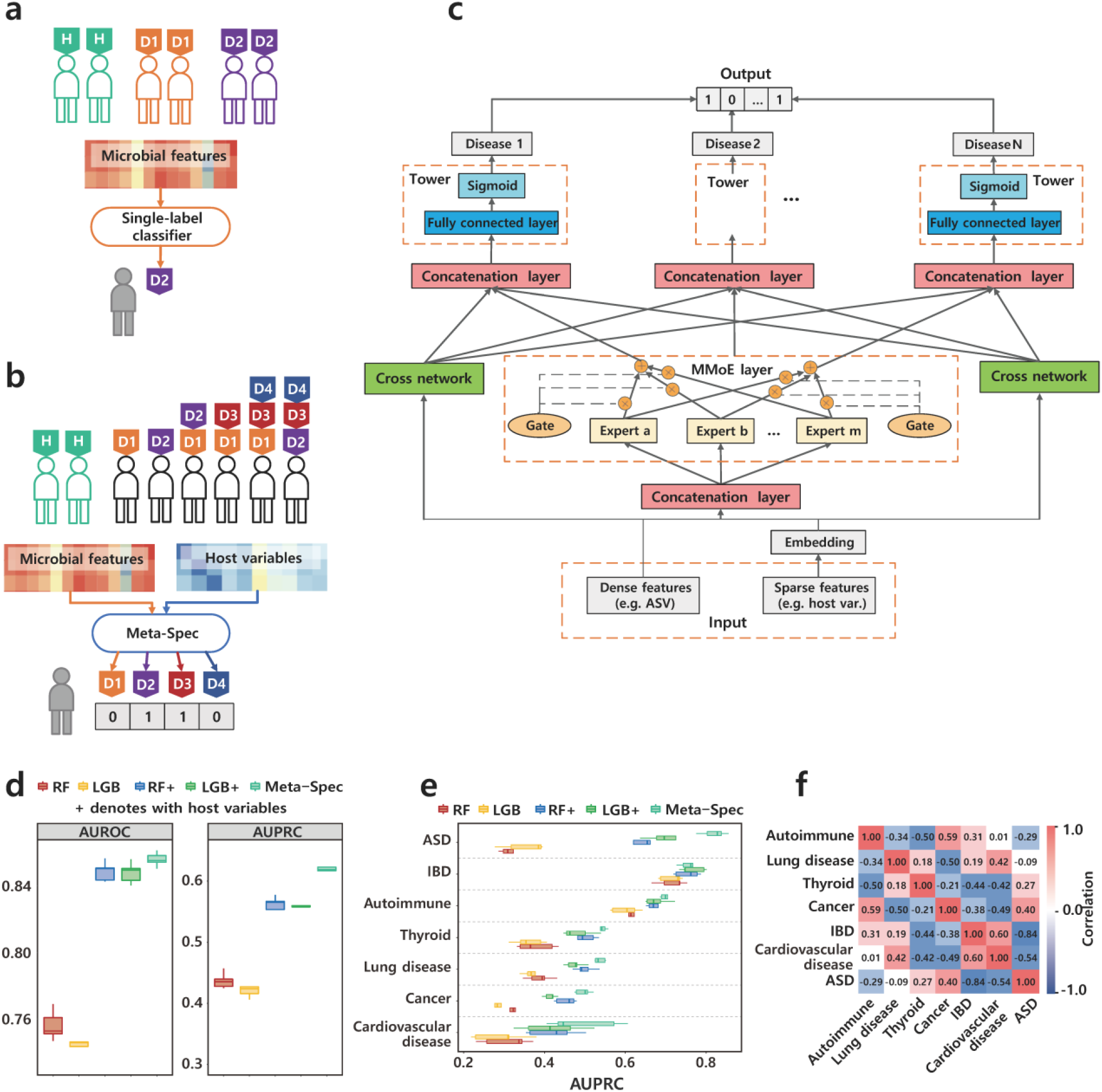
Multi-label disease classification. **(a)** In typical experiment design, each subject only has a single status label. (**b**) In actual cohort, a subject can have multiple diseases. **(c)** Deep learning framework of Meta-Spec. **(d)** AUROC and AUPRC of Dataset 1. (**e**) AUPRC of each disease of Dataset 1. (**f**) Disease correlations of Dataset 1.

On the other hand, the lifestyle and physiological variables of human hosts have strong connections to various diseases, which can also interfere with the recognition of microbiome-based status. Age, for instance, is one of the major risk factors for cardiovascular disease [10], and is also associated with Crohn’s disease [11]; Moreover, body shape and BMI can act as predictors for metabolic syndrome [12] and type 2 diabetes [13]. These host variables provide crucial information that can aid in disease prediction and diagnosis. Nevertheless, many existing ML classifiers focus solely on analyzing microbiome features, such as microbial diversity, abundance, and composition, while overlooking the potential of metadata in disease screening. Although these microbial features are important in disease screening and prediction, they are not the only factors that contribute to disease risk.

Here we present Meta-Spec, an explainable deep learning method for multi-label disease classification using microbiome data. Meta-Spec is based on a multi-gate mixture-of-experts (MMoE) model [14] and cross networks [15], which is capable to detect multiple diseases simultaneously by integrating genotype data (microbiome features derived from sequences) and phenotype data (host variables). In addition, unlike other neural network-based methods that lack interpretability, Meta-Spec can quantify the importance of confounding factors in describing each disease by their relative contribution in status classification. Experiments on multiple cohorts showed that the performance of our method was superior to widely used ML classification strategies in comorbidities screening and disease correlation catching, while also provide insights into the underlying mechanisms of each specific disease.

## Results

### Multi-label disease classification based on multi-task deep learning

The framework of Meta-Spec for multi-label classification is illustrated in **Figure 1c**. During model training, richness of microbial features (e.g., taxa, amplicon sequence variants (ASVs), operational taxonomy units (OTU), functional gene families, etc.) are treated as dense features, while host variables (e.g., physiological characteristics and lifestyle habits) are transformed into high dimensional embedding vectors, representing sparse categorical features (details provided in the **Methods and Materials** section). Then the concatenation layer merges dense features and embedding vectors, followed by an MMoE layer for learning disease associations, and two cross networks for capturing microbial and host variable interactions, respectively. Finally, the tower network calculates each disease’s probability by combining the outputs of the MMoE and cross networks. Thus, with a new microbiome and corresponding metadata, Meta-Spec classifier generates a binary array to summarize the prediction results, where each bit represents a particular disease’s presence (**Figure 1b** and **Figure 1c**).

### Deep neural network largely improves the multi-label disease classification

We evaluated the efficacy of Meta-Spec in multi-label disease screening using Dataset 1 (**Table 1**; see **Methods and Materials** for details), which was produced by the American Gut Project [8]. To minimize the impact of country information on gut microbiome [16], here we only employed the US cohort. This dataset contained 5,308 subjects, including 3,767 patients and 1,541 healthy controls. We filtered out invasive inspection information (e.g. blood) and only kept questionnaire-based host variables. Each patient in this dataset was diagnosed with at least one of the 7 target diseases of autoimmune disease, lung disease, thyroid disorder, cancer, inflammatory bowel disease (IBD), cardiovascular disease, and autism spectrum disorder (ASD). Although previous studies have demonstrated the association between gut microbiome and these target diseases [17-19], screening for status can be significantly affected by the combined influence of multiple diseases. We utilized 5-fold cross-validation to perform multi-label disease classification using Meta-Spec and other machine learning (ML) classifiers, including random forest (RF) that has been commonly used in microbiome studies [5, 20]) and light gradient boosting machine (LGB), which is the latest gradient boosting method developed by Microsoft [21, 22]). Performance was evaluated using the regular *AUROC* (area under the receiver operating characteristic curve). In addition, since the number of samples were highly unbalanced among hosts and labels, we also compared the performance using *AUPRC* (area under the precision recall curve), which is sensitive to unbalanced datasets.

**Table 1.**
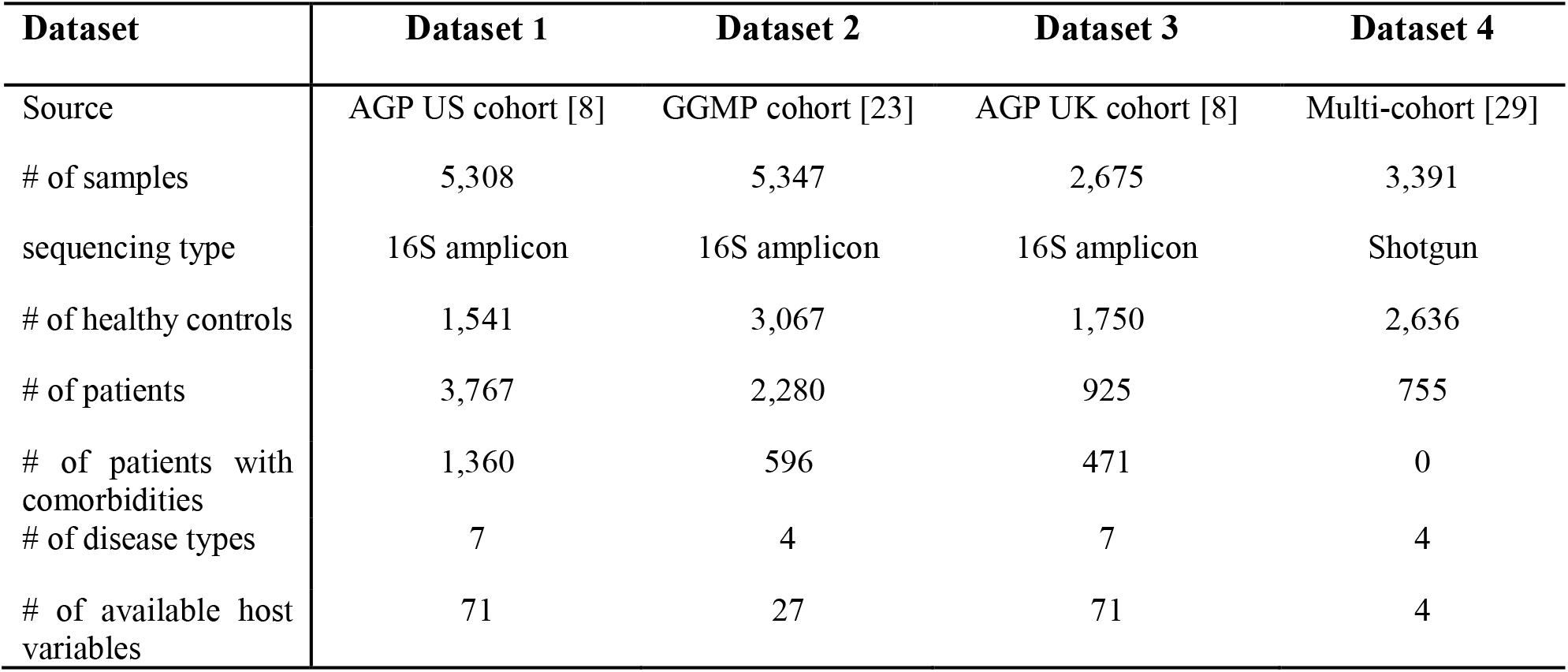
Dataset information.

| Dataset | Dataset 1 | Dataset 2 | Dataset 3 | Dataset 4 |
| --- | --- | --- | --- | --- |
| Source | AGP US cohort [8] | GGMP cohort [23] | AGP UK cohort [8] | Multi-cohort [29] |
| # of samples | 5,308 | 5,347 | 2,675 | 3,391 |
| sequencing type | 16S amplicon | 16S amplicon | 16S amplicon | Shotgun |
| # of healthy controls | 1,541 | 3,067 | 1,750 | 2,636 |
| # of patients | 3,767 | 2,280 | 925 | 755 |
| # of patients with comorbidities | 1,360 | 596 | 471 | 0 |
| # of disease types | 7 | 4 | 7 | 4 |
| # of available host variables | 71 | 27 | 71 | 4 |

The results presented in **Figure 1d** demonstrated that using only the microbiome ASVs, both RF and LGB achieved low AUROC values. This was likely due to the confounding effects of host factors or multi-disease interactions. When metadata was included through one-hot coding, the performances of both models were substantially increased, highlighting the significant role of host variables in disease detection. Meanwhile, it should be noted that the RF and LGB still exhibited shortage in the detection of ASD and Thyroid diseases indicated by detailed AUPRC values, as seen in **Figure 1e**. The best overall performance of both AUROC and AUPRC was achieved by Meta-Spec, as shown in **Figure 1d**, which outperformed all other models by a significant margin. This can be attributed to the deep neural network of Meta-Spec, which was able to capture disease associations during model training, as shown in **Figure 1f**. For example, the network identified a positive correlation (PCCs = 0.60; Pearson Correlation Coefficient) between IBD and cardiovascular disease, as well as a negative correlation (PCCs = -0.84) between IBD and ASD in Dataset 1.

We also validated Meta-Spec by Dataset 2 (**Table 1**) collected from the Guangdong Gut Microbiome Project [23], which consists of 5,347 subjects (**Table 1**; refer to **Methods and Materials** for details). Patients were diagnosed with at least one target disease of metabolic syndrome, gastritis, type 2 diabetes mellitus (T2DM) and gout. For Dataset 2, results were in the same trend as those of AGP US cohort, i.e. low AUPRC only by OTUs, then significantly raised with additional metadata in classification (**Figure S1**), verifying the superiority of multi-label classification strategy in Meta-Spec (**Figure S2**). In addition, Meta-Spec also revealed a positive connection between gastritis and gout in Dataset 2 (PCCs = 0.43; **Figure S3**).

### Disease-correlation are crucial for comorbidity detection

On the other hand, comorbidities are prevalent in actual cohorts, as evidenced by the fact that out of 3,767 patients in Dataset 1 cohort, 1,360 were identified as having two or more diseases. These comorbidities often go unnoticed by regular classification strategies, which is why we conducted further measurements to assess the ability of different methods in detecting them. To accomplish this, we divided patients of Dataset 1 and Dataset 2 into two groups (as shown in **Figure 2a**): the single-disease group with only the target disease and the comorbidity group with additional disease(s). The comorbidity detection results were based on multi-label classification, which took into account the performance on both the target disease and the comorbidities.

**Figure 2.**
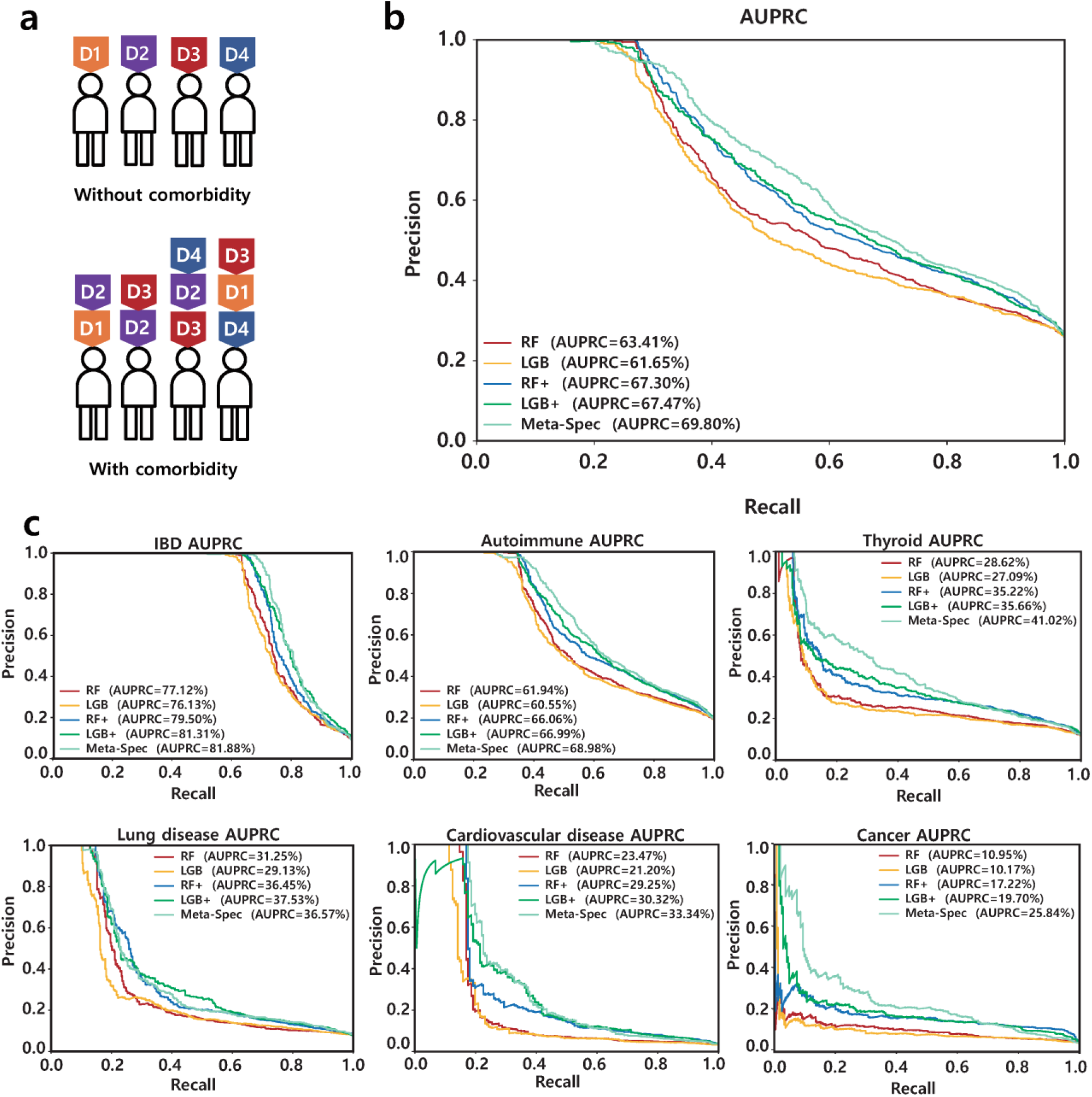
Performance of comorbidities detection. **(a)** Patients were divided by comorbidity status (**b**) Overall AUPRC of Dataset 1. **(c)** detailed AUPRC of Dataset 1.

As illustrated in **Figure 2b**, using microbiome structure and host variables for training, Meta-Spec achieved significantly higher AUPRC than both LGB and RF (**Figure 2b**). Such advantage mainly benefited from disease associations inferred by MMoE gates (**Figure 1f**), as well as the combination of multiple host variables by the cross network in Meta-Spec. On contrary, LGB and RF can miss comorbidities when focusing solely on target diseases like IBD, autoimmune and thyroid (**Figure 2c**).

### Feature selection and variable refinement for multi-label disease screening

To tackle the shortage of model interpretability of most neural networks, Meta-Spec is able to quantify the contribution of microbial members and confounding factors in recognizing disease patterns using a Meta-Spec Importance value (MSI; refer to **Methods and Materials** for details). Sorted by MSI, host variables were the dominant features for disease screening on Dataset 1, although their values varied among different diseases ((**Figure 3a**; **Figure S4**). Taking cardiovascular disease as an example (**Figure 3a**), age was the most important feature for cardiovascular disease detection. Specifically, old people, artificial sweeteners and people who have constipation were more susceptible to cardiovascular disease [10, 24, 25]. The highly ranked MSI values of host phenotypes also elucidate their effects to improve the performance of RF and LGB for cardiovascular disease classification (**Figure 1e**). Besides, some features analyzed from microbiome sequences also helped in distinguishing the disease, including ASV 1 (*Escherichia_Shigella*), ASV 28 (*Bacteroides*) that have been reported [26, 27], yet they received weaker importance in Meta-Spec model.

**Figure 3.**
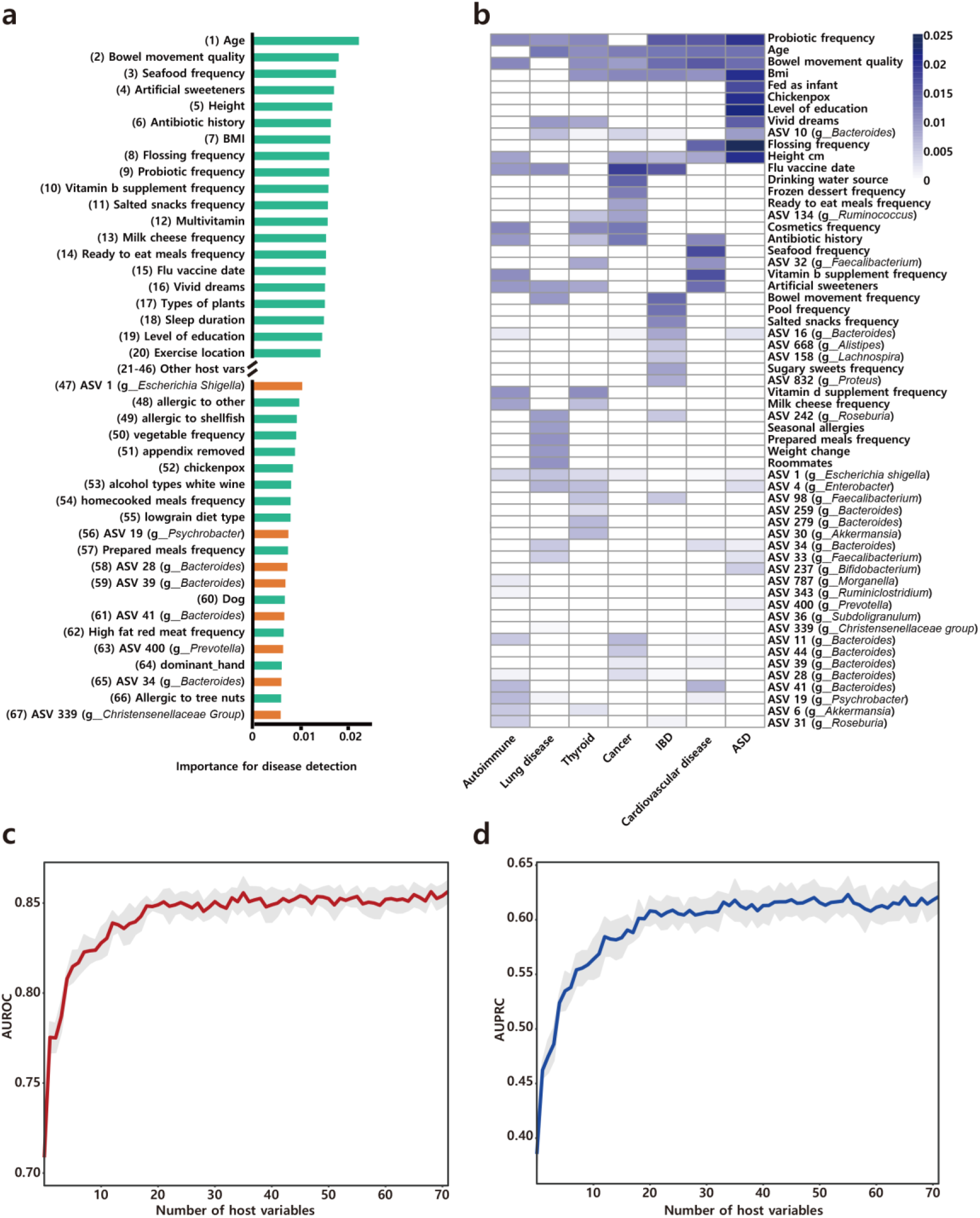
Meta-Spec Importance and host variable refinement. **(a)** Top host variables and microbial members for cardiovascular disease sorted by MSI. Numbers are their actual ranks. **(b)** Distribution of most importance variables among diseases in Dataset 1. **(c)** Change of AUROC of Meta-Spec with different number of host variables. **(d)** Change of AUPRC of Meta-Spec with different number of host variables.

By analyzing the mostly contributed features (top 20% MSI of host variables and microorganisms) of each disease, we observed that over 80% of them were shared by at least two diseases on the two datasets (**Figure 3b** and **Figure S5**; **Table 2**). Such non-specific associations among gut microbiota, host variables and diseases [28] further explained the limitation of bio-marker-based and regular ML-based strategies in comorbidities detection. Among them, a few factors were highly-ranked in most diseases, such as BMI, bowel movement quality, probiotic frequency, ASV 1 (*Escherichia_Shigella*) and ASV 10 (*Bacteroides*) of Dataset 1 (**Figure 3b**); and districts, medication, OTU 4425571(*Escherichia_Shigella*) and OTU 136025 (*Ralstonia*) of Dataset 2 (**Figure S5**). On the other hand, we also found the distinctive features that were only sensitive to a single disease. For instance, seafood frequency was strongly associated with cardiovascular disease, as well as OTU 4478762 (*Lacticigenium*) and BMI contributed to Metabolic syndrome detection. Nevertheless, although these features were important for disease detection, none of them can work as the disease indicator solely. For example, as universal markers, age, waist size, medication and ASV 1 only achieved the AUPRC of 0.300, 0.248, 0.173 and 0.162 in status predicting (**Figure S6**), largely lower than then overall performance. Hence proper classification methods and models are essential to fully explore the significance of important factors.

**Table 2.** Number of features among diseases.

|  | Dataset 1 | Dataset 2 |
| --- | --- | --- |
| # of total features | 1,154 | 476 |
| # of mostly contributed features (union set of all diseases) | 372 | 128 |
| # of commonly shared features | 150 | 84 |
| # of specific features for a disease | 66 | 24 |

After the preliminary artificial curation of the original metadata, there were still 71 host variables kept in model training and validation for Dataset 1. Since too many items in questionnaires can introduce difficulties in real applications, a feature selection procedure was performed to assess the effect of metadata amount in classification, thus reducing the number of enrolled host variables. For Dataset 1, we first sorted all the host variables by their mean MSI over all target diseases. Then, less important host variables were gradually eliminated to repeat the model training and the corresponding validation by Meta-Spec. The performance curves (**Figure 3c** and **Figure 3d**) described the linkages between the performance of multi-label classification and the number of enrolled host variables. We noticed that when taking only 20 host variables, Meta-Spec can still provide promising multi-disease screening. Such effort is crucial in actual application scenarios for Meta-Spec enhanced the potential of gut microbiomes in multiple diseases screening with a little easy-collected information from hosts.

### Meta-Spec with hybrid model expands the application of microbiome across geographical locations

Geography has been shown to have a significant impact on the variations in the human microbiome [16, 23]. Making a data model built from the local cohort the most suitable option for microbiome-based detection, but limited training samples can pose a challenge to this approach. In such cases, adopting well-validated models from other locations becomes a more practical option. However, the applicability and compatibility of the model need to be taken into consideration. To study the cross-cohort multi-label classification of Meta-Spec, we conducted a 5-fold cross-validation on the UK cohort of the AGP dataset (**Table 1**, Dataset 3). In each fold, we used 1/5 of the UK samples as the test set, and two different sets for training (**Figure 4a**) of *a*. the local set (comprising other 4/5 UK samples); and *b*. the hybrid set (comprising other 4/5 UK samples and US samples). Moreover, for both of the two training sets, we varied the amount of UK training samples from 10% to 100% to simulate the lack of local samples for modeling.

**Figure 4.**
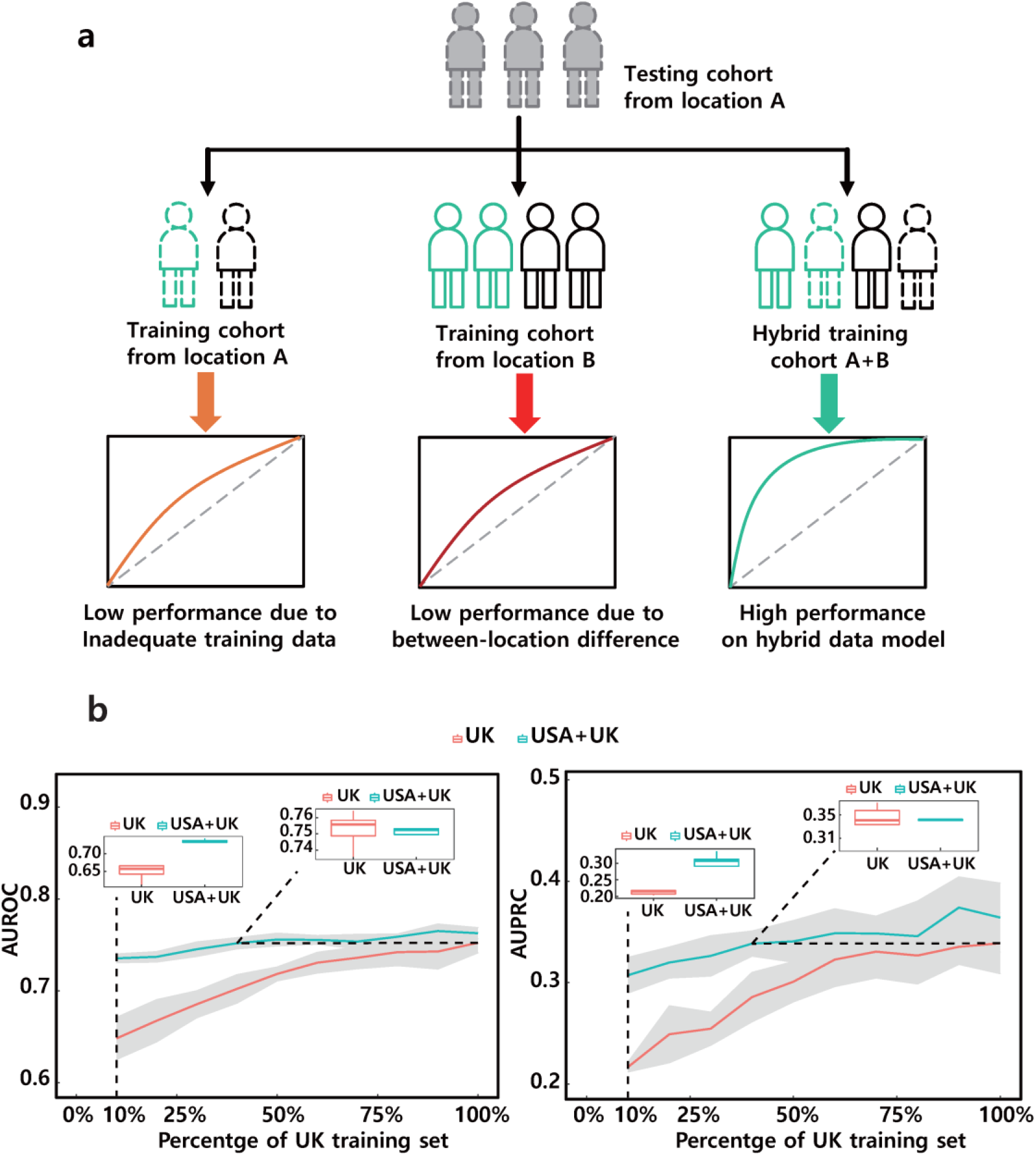
Performance comparisons of cross-cohort validation. **(a)** Hybrid modeling for cross-location detection. **(b)** Overall AUROC and AUPRC trend with the increase of UK training samples in hybrid modeling

As expected, the UK local cohort with limited learning data (only 10% of the training set) exhibited poor classification performance (as shown in **Figure 4b**). However, when combined with microbiome features and host variables from the US cohort, the detrimental effect of data scarcity was mitigated, resulting in a significant improvement in cross-cohort classification results. By introducing more UK samples into both sets, we found that the hybrid set consistently outperformed the UK local cohort in terms of AUPRC and AUROC on the performance curve. Even with only 40% UK samples, the hybrid set achieved similar levels of performance as those obtained by utilizing the entire UK local set. Furthermore, the performance of the local set, comprising 100% UK samples, can be further optimized and enhanced by incorporating cross-cohort data. Thus, despite the challenges posed by inter-location variations in the gut microbiome, host variable embedding and multi-cohort model integration offer significant technical advantages in bridging this gap in status classification.

### Applicability of Meta-Spec on regular single-label classification

To further verify the applicability of Meta-spec on widely studied single-label classification, we also employed the Dataset 4 with 3,391 metagenomes from multiple cohorts (**Table 1**; refer to **Methods and Materials** for details) [29]. Here each patient was labeled by a single disease from 4 categories of acute cerebral vascular disease (ACVD), colorectal cancer (CRC), Crohn’s disease and type 2 diabetes (T2D). With this dataset, two types of single-label classification test were performed using 5-fold cross validation: *i*) binary classification that only distinguishes disease samples from healthy controls. We also calculated the Gut Microbiome Health Index (GHMI) [29] for each sample to predict the likelihood of disease. As shown in **Figure 5a**, RF and LGB exhibited higher AUROCs than GMHI in binary status classification using microbiome taxonomy features. When further embedding 4 available host variables including gender, age, BMI and geographical region in model training, the average AUROC was improved up to 0.97. *ii*) multi-category classification that specifies the detailed disease type. **Figure 5b** suggested the prediction models were sharply enhanced by additional host information in the overall kappa coefficient. Therefore, compared to regular machine learning methods, Meta-Spec can also offer optimal performances in single-label disease screening.

**Figure 5.**
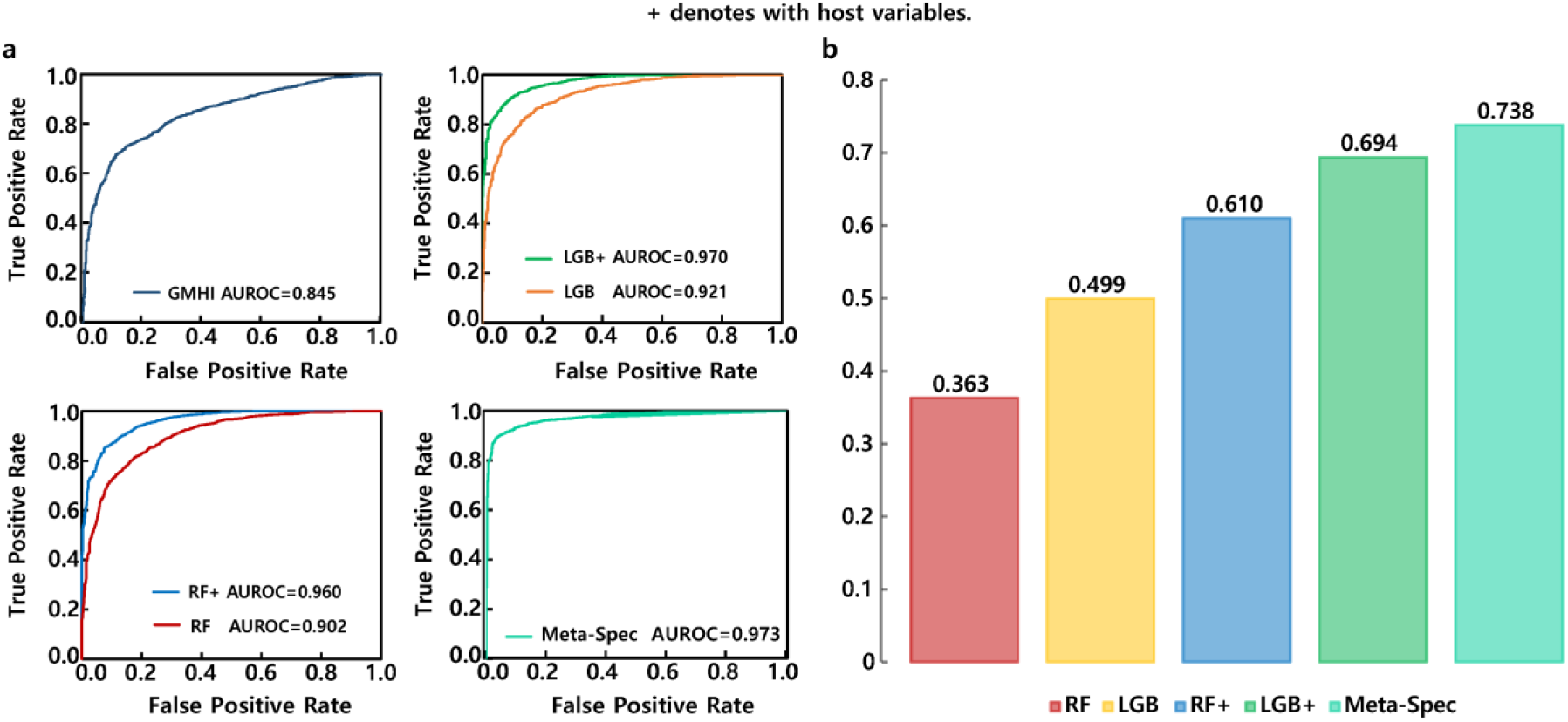
Performance comparisons on multi-class dataset. **(a)** ROC curve on healthy status detection. **(b)** Performance comparisons on multi-class prediction by Kappa.

## Conclusion and discussion

For years, researchers have been fascinated by the potential link between the human microbiome and various diseases. By studying microbial features, scientists hope to predict changes in human health status. However, this is a complex challenge, as disease interactions and variations in host lifestyles can interfere with the human microbiota. While the gut microbiota has been shown to play a critical role in human health, it’s important to consider the impact of host information on disease screening. Although these variables have been considered in experimental design, cohort recruitment, multi-variant statistics, and effect size measurement, they have not yet been included in microbiome-based modeling. By incorporating easily-collected host phenotype data, such as diet, BMI, and age, disease screening models can be significantly improved in terms of sensitivity and precision compared to those that rely solely on microbial features.

On the other hand, machine learning has been increasingly used to develop prediction models. However, many training processes still rely on traditional ML techniques, such as SVM and RF, which do not take advantage of the latest developments in ML or DL. By using multiple datasets and cohorts, we have demonstrated the benefits of a cutting-edge deep neural network for multi-disease classification of biological data with inherent complexity. Furthermore, by treating each label as a single task, our Meta-Spec approach can be quickly and easily updated to accommodate additional diseases with only minor modifications, while regular ML models require significant reconstruction. These efforts represent an important step towards understanding the underlying properties of unknown microbiomes.

## Methods and Materials

### Host variable embedding and deep learning framework of Meta-Spec

A microbiome sample can be demoted as a vector *x* = (*x*_1_, 舰, *x*_*h*_, *x*_*h*+1_, 舰, *x*_*d*_), where the first *h* features are sparse features (host variables) and the last (*d* - *h*) features are dense features (microbial members). Since sparse features is much less than dense features (*h* << *d* – *h*), the effect of host variables can be diluted in modeling due to the imbalanced feature number. To tackle this problem, in Meta-Spec, we encode each sparse feature as an *m*-dimensional embedding vector (*m* was set as 128), and all embedding vectors are then re-integrated with dense features into a (*d* - *h* + *m*\**h*) dimension vector *c* ∈ ℝ^(*d*−*h*)+*m*\**h*^.

This high-dimensional vector *c* is used to train multiple experts as both features and parameters. As two-layer DNNs, these experts can learn associations among diseases by sharing parameters across all tasks and the output of the *l*_*th*_ expert is denoted by *f*_*l*_(*c*). Besides multiple experts, MMoE layer sets a gate for each disease to generate weights and assemble outputs of multiple experts. 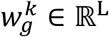 is the weight vector of the *k*_th_ gate, and the MMoE output for the *k*_th_ tower network *T*^*k*^(*c*) is obtained by weighting the output of experts as *equation 1*:

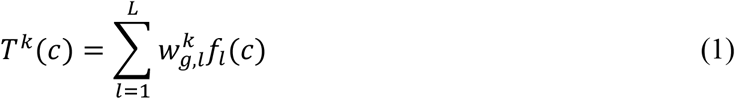

To capture microbial interactions, a dense cross network is constructed for microbes by *equation 2*:

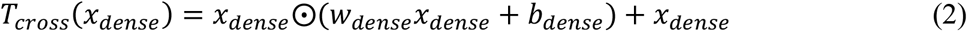

where *x*_*dense*_ ∈ *R*^*d*−*h*^ is a combination of all dense features, *w*^*dense*^ ∈ *R*^(*d*−*h*)×(*d*−*h*)^ is a weight matrix, *b*_*dense*_ ∈ *R*^*d*−*h*^ is a bias vector and ⨀ represents Hadamard product. Similarly, we also develop a cross network to learn host variable interactions by *equation 3*:

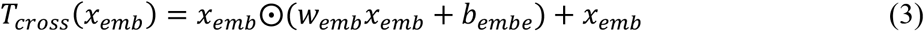

where *x*_*emb*_ ∈ *R*^*h*×*m*^ is a combination of all sparse features, *w*_*emb*_ ∈ *R*^(*h*×*m*)×(*h*×*m*)^ is a weight matrix and *b*_*emb*_ ∈ *R*^*h*×*m*^ is a bias vector.

For each tower network, the outputs of the two cross networks are concatenated with the corresponding MMoE output as its input. Each tower network consists of a fully connected layer with a sigmoid to output the final predictions. Additionally, an automatic weighted loss function [30] is employed to combine multi objective losses by *equation 4*:

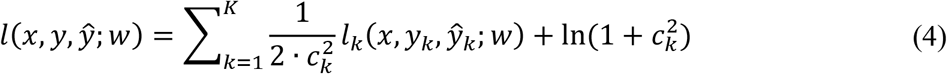

where *c*_*k*_ is the trainable weight of the *k*_th_ task, *w*_*k*_ is the network parameters and *l*_*k*_(*x, y*_*k*_, *ŷ*_*k*_; *w*_*k*_) is the loss function of the *k*_th_ task.

During model construction, different from traditional ML approaches that treat input features as input constants, Meta-Spec continuously updates the embedding vectors by iterations. Meanwhile, the two cross networks can also capture microbial interactions and host variable interactions respectively. Therefore, Meta-Spec can not only learn associations among diseases but also take advantage of sparse features, which make it outperform traditional machine learning methods.

### Meta-Spec Importance calculation

To rank and quantify the contribution of microbial features and host variables in disease screening, we defined a Meta-Spec Importance (MSI) based on SHAP [31] derived from game theory [32]. With a Meta-Spec model and a test dataset, the MSI value explains the proportion of a feature’s contribution to the prediction. More specifically, for a feature *i*, first we parse out its relative contribution *C*_*i*_ by *equation 5*:

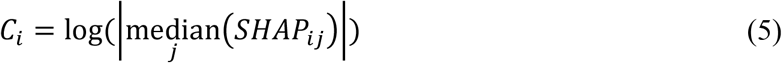

where *SHAP*_*ij*_ denotes the SHAP value of the *ith* feature of the *jth* sample in the test set. Then the MSI is generated by the normalization of contribution *C*_*i*_ in *equation 6* and *equation 7*:

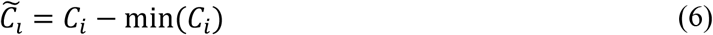

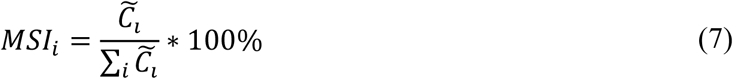

In this way, for a specific disease, the total sum MSI of all features is normalized as 100%.

### Performance evaluation and comparison

Here we also used RF and LGB to build regular binary classifiers for comparison. For multi-label classification, regular ML models were trained from the original vector *x* (refer to section **4.1**), and organized by binary relevance that decomposed the task into several independent binary classifiers (one per label). According to Statnikov et al.’s work [33], parameters for each model were tuned as **Table 3**. We applied nested repeated 5-fold cross-validation in the test procedure, which selected 80% of the data as the training set and 20% of the data as the test set.

**Table 3.** parameters of model tuning.

| Method | Parameter | Value |
| --- | --- | --- |
| Random Forest | n_estimators (number of trees) | {300, 500, 1000} |
| Random Forest | max_features (number of features) | $\{0.5\sqrt{\# \text{ of ASV}}, \sqrt{\# \text{ of ASV}}, 2\sqrt{\# \text{ of ASV}}\}$ |
| LGB | n_estimators (number of boosted trees) | {300, 500, 1000} |
| LGB | max_depth (maximum tree depth) | {4, 6, 8} |
| Meta-Spec | num_experts (number of experts) | {5, 7, 9} |
| Meta-Spec | dnn_hidden_units (number of hidden units) | $\{(256, 128), (128, 256), (128, 128), (256, 256)\}$ |

In each fold, *AUPRC, F1-macro* and *AUROC* were calculated to evaluate the performance. *AUROC* is the area under the ROC curve, while *AUPRC* stand for the area under the Precision-Recall curve. And the average *AUROC* and the average *AUPRC* across all diseases were treated as the overall *AUROC* and the overall *AUPRC*. Additionally, *F1-macro* averages the *F1* score on the prediction of different diseases by the following equations:

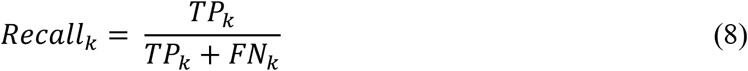

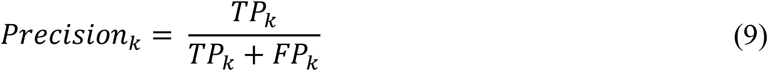

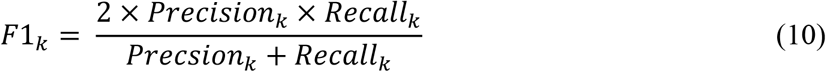

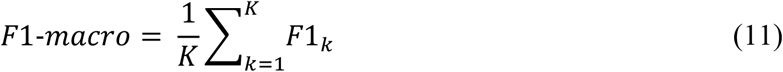

where *TP*_*k*_, *FP*_*k*_, *FN*_*k*_, *Recall*_*k*_, *Precision*_*k*_ and *F*1_*k*_ represent true positive, false positive, false negative, recall, precision and F1 for detecting the *k*_th_ disease.

### Microbiome datasets and pre-process

The brief information of all datasets is summarized in **Table 1**. Dataset 1 and Dataset 3 were collected from the American Gut Project (AGP) cohort [8]. ASVs of 16S rRNA gene amplicons and metadata of each subject were directed downloaded from Qiita (study ID:10317) [34]. The taxonomy of ASVs was then annotated by Greengenes 13-8 database [35] using Parallel-Meta Suite [36]. A subject was treated either as a patient if recorded as ‘Diagnosed by a medical professional (doctor, physician assistant)’ for a specified disease in the metadata, or as healthy if marked as ‘I do not have this condition’ for all diseases. To eliminate the sparsity of ASVs, we performed a distribution-free independence test based on the mean variance index [37, 38], and selected 1,168 ASVs that were relevant to healthy status for disease detection. Dataset 2 was collect from the Guangdong Gut Microbiome Project (GGMP) [23]. The 16S rRNA gene amplicon sequences and metadata of each subject were obtained from EBI (ID: PRJEB18535), and OTUs were picked by GGMP pipeline (https://github.com/SMUJYYXB/GGMP-Regional-variations). We also employed the distribution-free independence test based on the mean variance index and selected 449 OTUs that were relevant to the target diseases. Dataset 4 was a cross-cohort dataset produced by 34 studies [29]. Species-level taxonomy of raw shotgun stool metagenomes was analyzed by MetaPhlAn 2 [39]. Additionally, in each dataset, the chi-square test was utilized to select host variables that associated with at least one target disease.

## Code and data availability

The software package of Meta-Spec is released at GitHub (https://github.com/qdu-bioinfo/meta-spec). Source data of datasets used in this work is summarized in **Table 1**.

## Acknowledgements

X.S. acknowledges the support of grant 2021YFF0704500 from National Key R&D Program of China, grant 32070086 from National Nature Science Foundation of China, grant from Taishan Scholar Youth Expert program and Youth Innovative Talents Program of Shandong Province of China. S.W. acknowledges the support of grant ZR2019PF012 from Shandong Provincial Natural Science Foundation of China.

## Author contributions

X.S. and S.W. conceived the idea. S.W. and Z.L. implemented the algorithm and codes. Z.L. and

J.X. performed the analysis. Y.C., Y.S., F.Z. and M.Z. collected the datasets. X.S., R.K. and S.H. wrote the manuscript. All contributed the proofread.

