## Supplementarl materials for "Deep learning and host variable embedding augment microbiome-based simultaneous detection of multiple diseases"

**Supplementary materials for "Deep learning and host variable embedding augment microbiome-based detection of multiple diseases simultaneously"**

**Supplementary Figures**

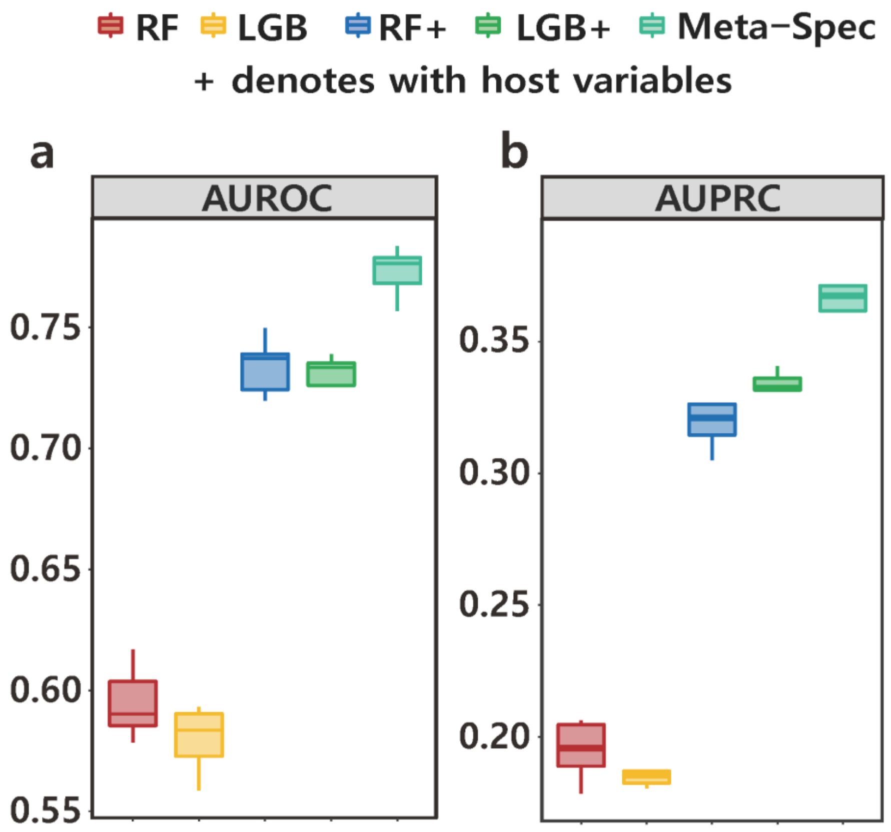


**Figure S1. Comparison of multi-label disease classification on Dataset 2 using on (a) AUROC and (b) AUPRC.** Source data is provided in Supplementary Data.

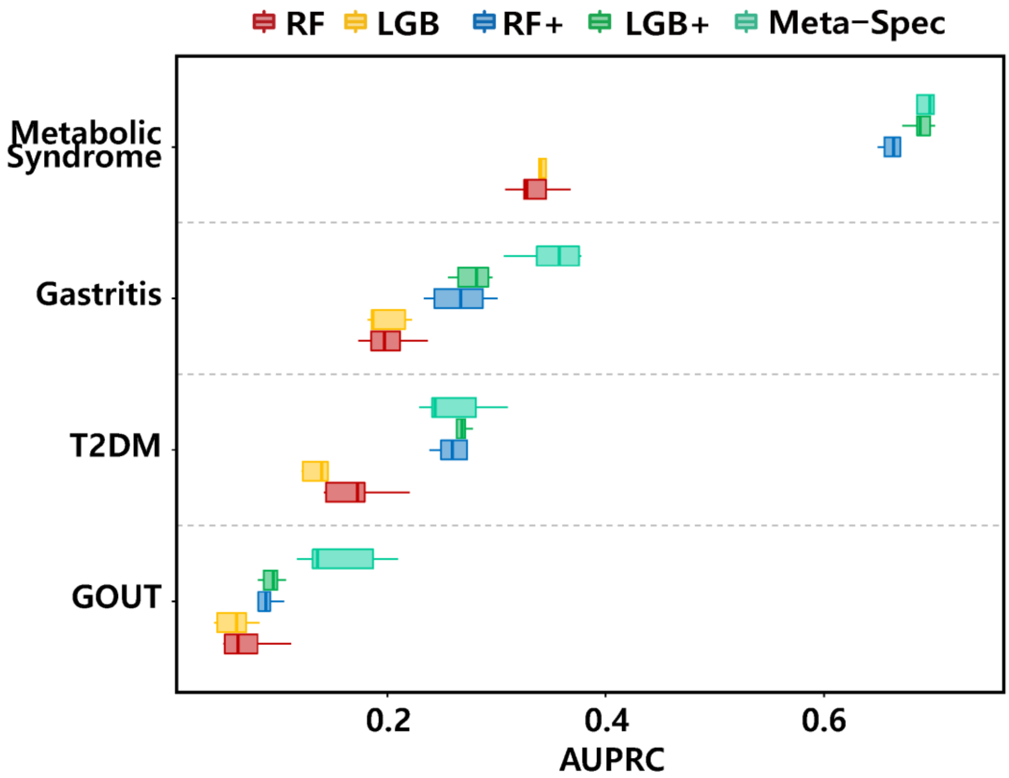


**Figure S2. AUPRC of each disease of Dataset 2.** Source data is provided in Supplementary Data.

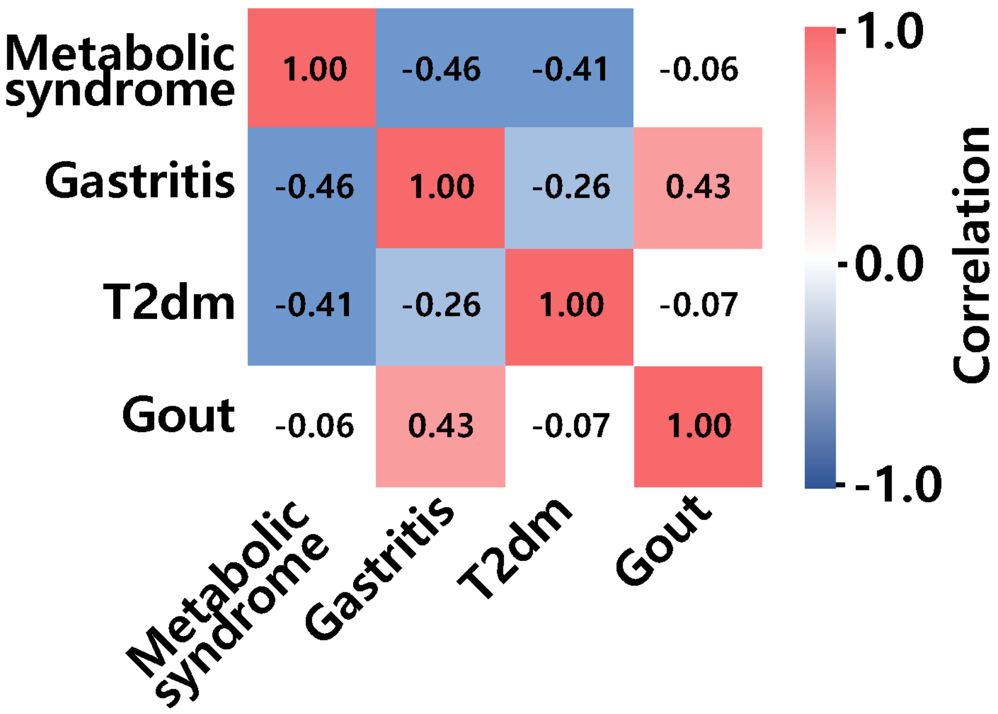


**Figure S3. Disease correlations of Dataset 2.** Source data is provided in Supplementary Data.

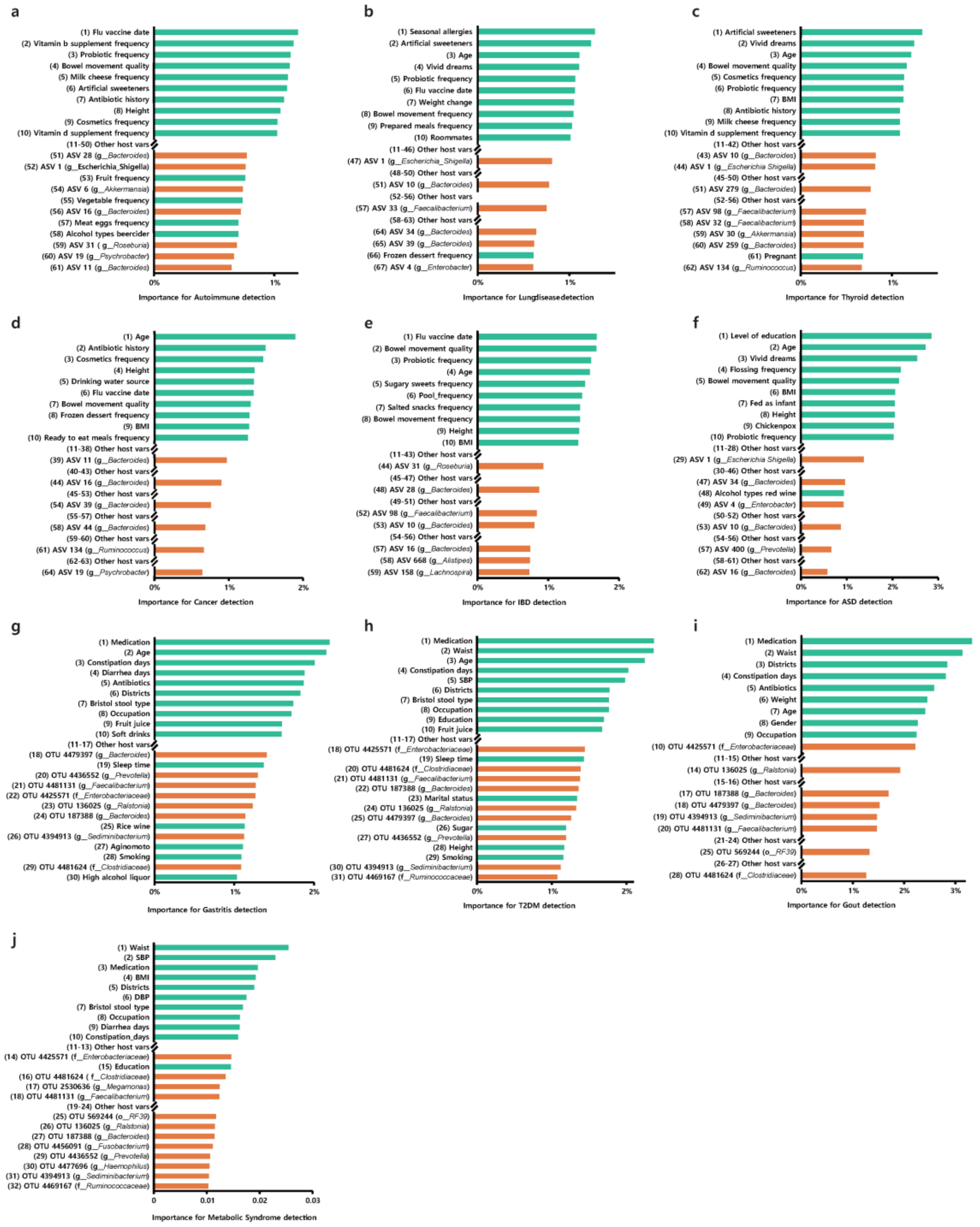


**Figure S4. Top host variables and top microbial members for disease detection sorted by importance of Meta-Spec Importance.** Numbers are their actual ranks. **(a)** Autoimmune. **(b)** Lung disease. **(c)** Thyroid. **(d)** Cancer. **(e)** IBD. **(f)** ASD. **(g)** Gastritis. **(h)** T2DM. **(i)** Gout. **(j)** Metabolic Syndrome. Source data is provided in Supplementary Data.

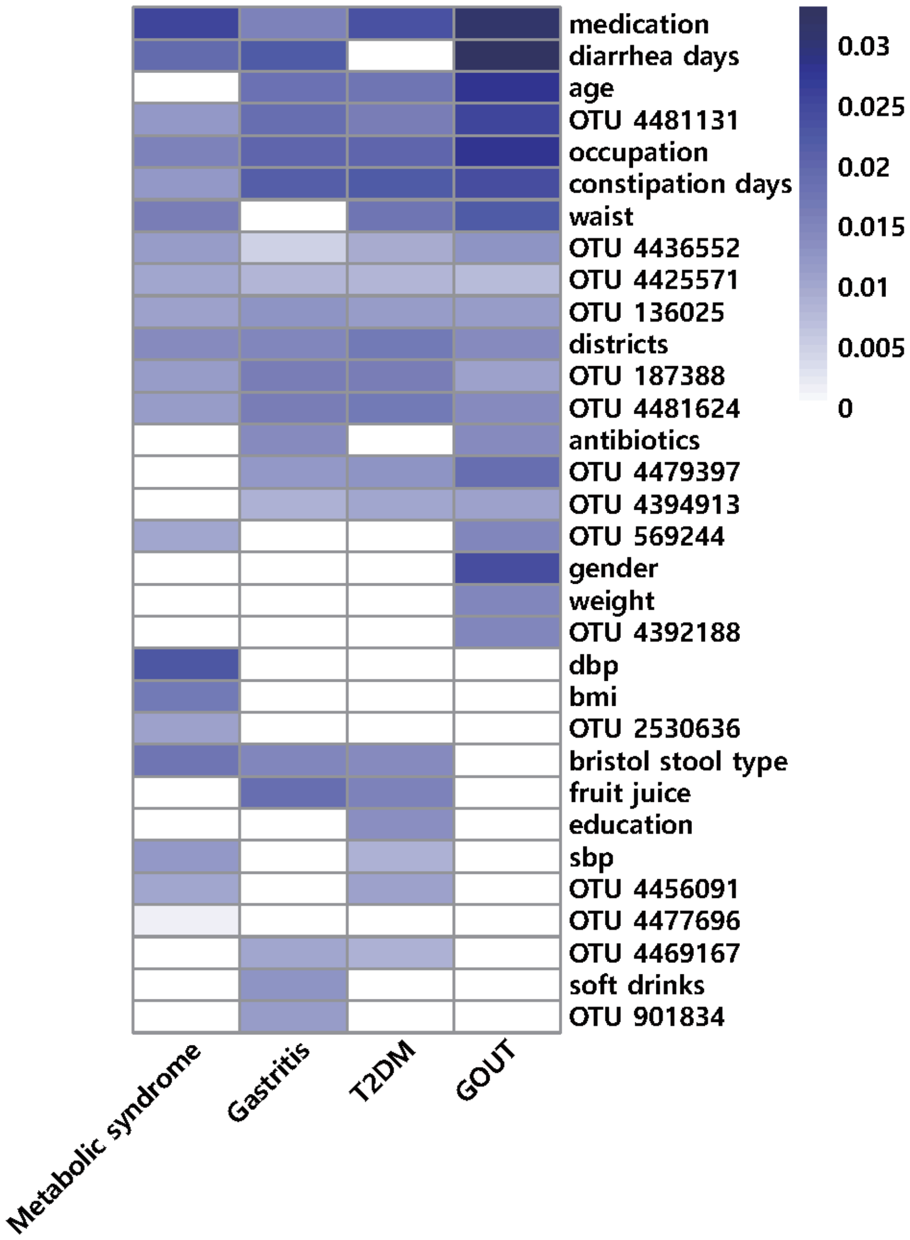


**Figure S5. Distribution of most importance variables among diseases in Dataset 2.**

Source data is provided in Supplementary Data.

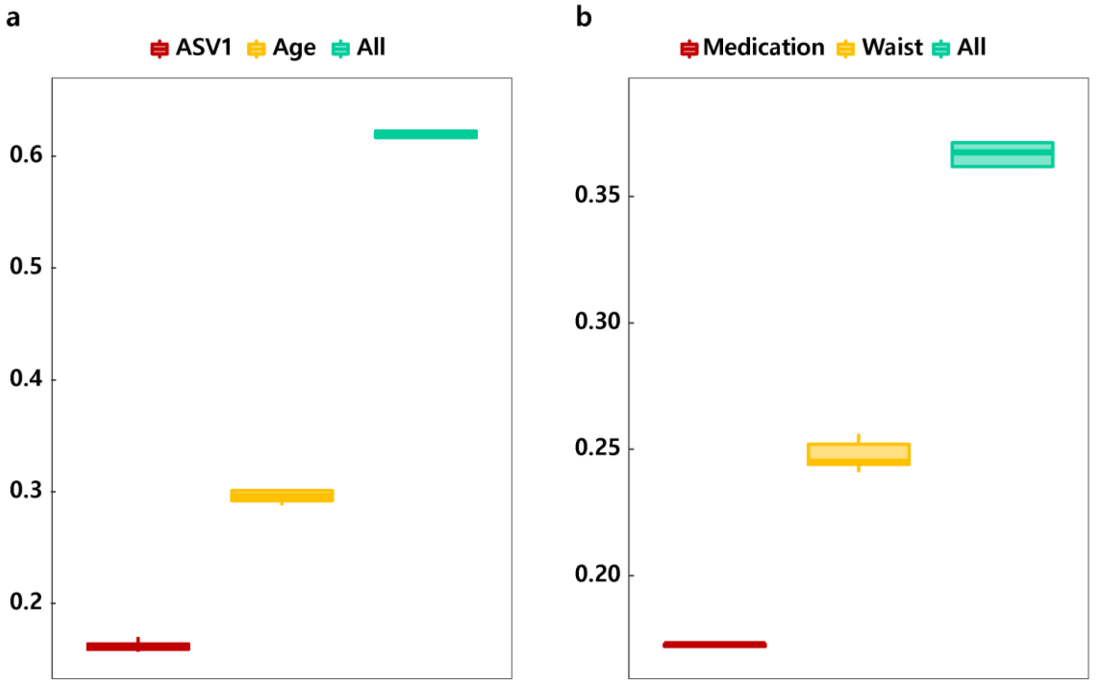


**Figure S6. AUPRC of multi-label disease classification using a single feature. (a)** ASV1 and age of Dataset 1 and **(b)** medication and waist size of Dataset 2. Source data is provided in Supplementary Data.
